# Transparent and flexible ECoG electrode arrays based on silver nanowire networks for neural recordings

**DOI:** 10.1101/2020.02.24.962878

**Authors:** Joana P. Neto, Adriana Costa, Joana Vaz Pinto, André Marques–Smith, Júlio Costa, Rodrigo Martins, Elvira Fortunato, Adam R. Kampff, Pedro Barquinha

## Abstract

This work explored hybrid films of silver nanowires (AgNWs) with indium-zinc oxide (IZO) for developing high-performance and low-cost electrocorticography (ECoG) electrodes. The transparent hybrid films achieved a sheet resistance of 6 Ω/sq enabling electrodes with 500 μm diameter to reach an impedance of 20 kΩ at 1 kHz and a charge storage capacity of 3.2 mC/cm^2^, an improvement in properties over IZO electrodes, whose performance is on par with the classical tin doped indium oxide (ITO). Characterization of light-induced artifacts was performed showing that light intensities <14 mW/mm^2^ elicit minimal electrical potential variation, which falls within the magnitude of baseline noise. The validation of electrodes in vivo was achieved by recording electrical neural activity from the surface of rat cortex under anaesthesia. Moreover, the presence of the hybrid films did not cause distortion of light during fluorescence microscopy. This study highlighted the capabilities of the hybrid structure of AgNWs with IZO, that can be fabricated with industrially-established processes at low cost, to be used for transparent ECoG electrodes, offering a new way to record neural electrical activity on a large and fast scale with direct visualization of neurons.

## INTRODUCTION

Brain function relies on communication between large populations of neurons across multiple brain areas. A full understanding how distributed populations of neurons coordinate their activity to generate behavior in animals will require tools capable of simultaneously monitoring neural activity on a large, precise and fast scale across multiple brain areas, ideally with the added ability to resolve genetically-distinct types of neurons^1,2^. In the past decades, technological advances have provided neuroscientist powerful tools for simultaneously monitoring the activity of thousands of neurons in vivo.

One such tool is functional calcium imaging (e.g., wide-field and two-photon imaging) where the activity of genetically-identified neurons can be detected optically using fluorescent calcium-sensitive dyes or proteins into the cell membranes of neurons. The primary strength of imaging approaches is their spatial resolution, allowing the relative locations, and the cell -type or connectivity of the neurons, to be obtained. The drawback of imaging-based approaches for recording neural activity is their poor temporal resolution (rise times of ~10-50 ms for single action potentials), thus they cannot give precise information about the timing of spiking in neurons relative to one another, or relative to behaviour^3,4^. *How can we monitor large-scale neuronal activity dynamics with high temporal resolution?* Extracellular recording (electrical recording) remains the only technique capable of measuring the activity of many neurons simultaneously with sub-millisecond precision^5^. ECoG is a form of extracellular recording and flexible ECoG electrode arrays (EAs) can cover the cortical surface providing a view of large-scale activity of distributed populations of neurons with high temporal resolution and minimal brain damage. However, the recorded signal cannot reveal exact locations of active neurons, and can only provide cell-type information if used in combination with photostimulation^6^, which is laborious and low-throughput. Each electrode is sensitive to the activity of hundreds of neurons in its vicinity^7^.

Simultaneous recordings of neural electrical activity on a large and fast scale with direct visualization of neurons could provide insight into how the brain works and to better understand the nature of the electrophysiological recorded signal. This would have an impact on clinical and brain-machine research where ECoG is frequently used.

There have been promising attempts to build transparent ECoG electrodes using ITO and graphene. An array with 49 ITO electrodes (500 μm diameter) was tested in vivo^8^. Thunemann et al. have recently reported the use of graphene in ECoG electrodes^9^. Arrays with 16 graphene electrodes (100×100 μm separated by 300 μm) enabled simultaneous 2-photon microscopy, optogenetic stimulation, and cortical recordings. While graphene presents remarkable properties, its fabrication by reproducible and large-scale methods is still challenging. Regarding ITO, due to indium scarcity and brittleness, alternative materials are currently being studied for solar cells and displays. These include conductive polymers, carbon nanotubes and metallic nanostructures - metal nanowires, thin metal films and patterned metal grids^10^. Among these, metallic nanostructures are considered to be the best candidates because of their inherently high electrical conductivity, optical transparency, mechanical robustness and, cost-competitiveness^11^.

In this work we have fabricated transparent electrodes with AgNWs and IZO, which enables a flexible, transparent and conductive hybrid structure that is stable under physiological conditions. Metal nanowires network maintain the excellent optical and electrical properties of patterned metal films^12,13^ adding simpler and lower cost manufacturing, turned available with low-cost solution-based deposition techniques. However, despite the advantages of AgNWs, some of their intrinsic drawbacks, such as silver oxidation and poor adhesion between nanowires and substrate, limit their practical application in ECoG electrodes. IZO, unlike ITO, can be processed at a low temperature and works as corrosion inhibitor to prevent the slow degradation of AgNWs film conductivity over time, whilst improving nanowires adhesion^14,15^. Therefore, the motivation for this work comes from the need to explore emerging materials to advance transparent ECoG electrodes, which are necessary to acquire large-scale neuronal activity with high spatio-temporal resolution.

## METHODS

### Fabrication and characterization of transparent and conductive films

Glass and silicon wafers were sonicated sequentially in acetone and isopropyl alcohol (IPA) for 15 minutes, rinsed with ultra-pure water, and finally dried with a stream of N_2_. IZO films (In_2_O_3_/ZnO, 89.3/10.7 wt%, 3″ ceramic targets from SCM, Inc.) were produced in a homemade RF magnetron sputtering system at RT^16,17^. The oxygen partial pressure and the deposition pressure (Ar + O_2_) were kept constant at 1.5 × 10^−5^ Pa and 1.0 × 10^−3^ Pa, respectively. The target– substrate distance and RF power were kept constant at 15 cm and 50 W, respectively. IZO films were deposited with thicknesses of 140 and 220 nm. Commercial AgNWs (115 nm diameter and 20-50 μm length) purchased from Sigma Aldrich were used with a concentration of 1.44 M. AgNWs were spray coated at 1 ml/min, directed by a stream of dry nitrogen gas at 7 Bar, onto clean glass substrates on a heated stage (150 ± 10 °C) using an ultrasonic spray head (Sonotek, 0.8 W), with a raster pattern design to maximize uniformity over the sample area (see in supplementary information Figure 1 the pattern of one cycle of spray deposition). To study the effect of network density on the optoelectronic properties, films with different number of cycles were produced. The control of network density was achieved by using a homemade in situ resistance monitor made using a 2.5 × 2.5 cm^2^ glass substrate, copper tape connected at the ohmmeter and silver paste to connect from copper to the glass substrate. On selected AgNW networks 140 nm thick IZO films were deposited to create the hybrid films.

**Figure 1.**
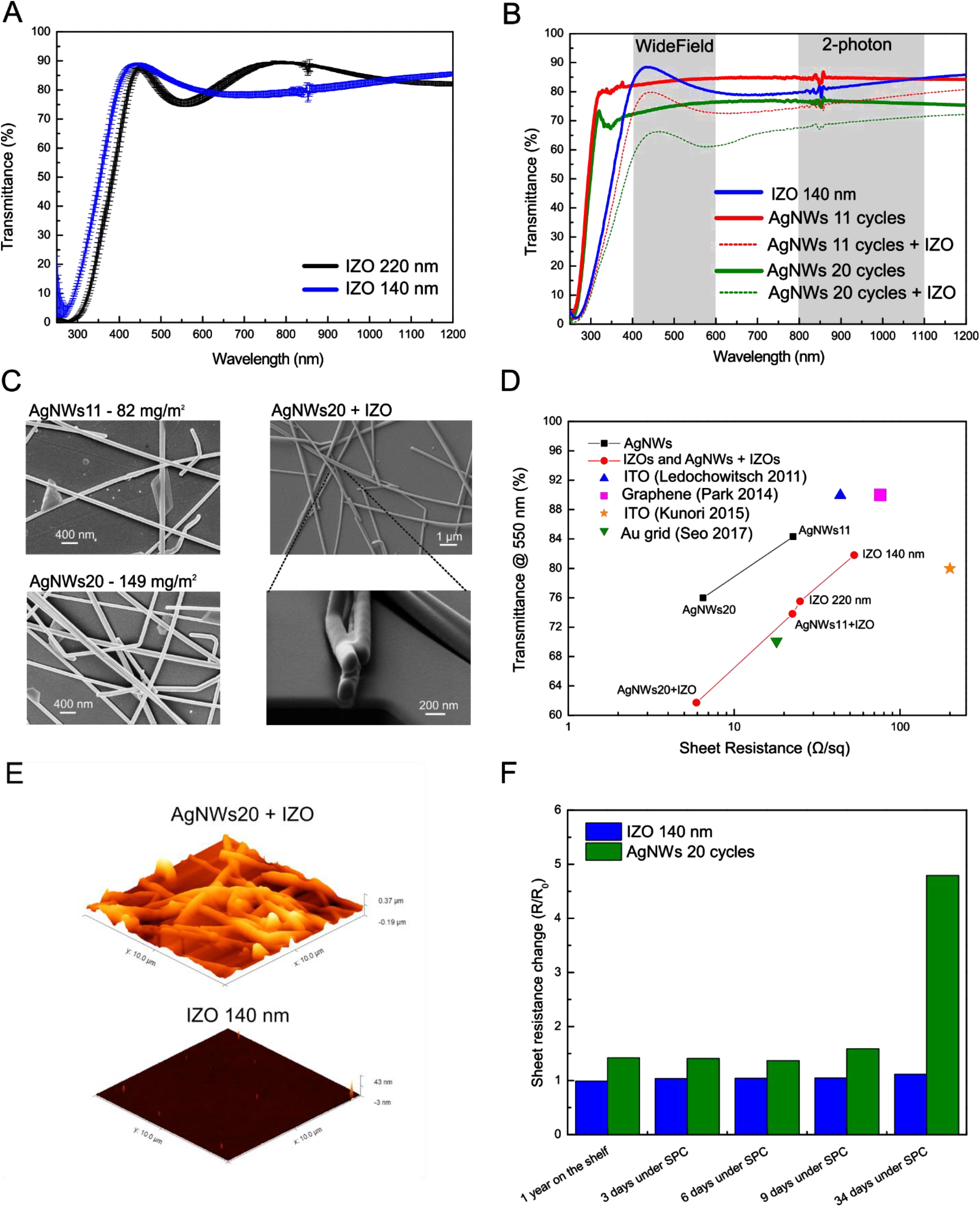
Film characterization. (a) Optical transmittance of IZO films with thickness of 140 and 220 nm; (b) Optical transmittance of AgNWs and hybrid films. The wavelength range used in widefield and two-photon microscopy is shown in grey shadow; (c) SEM images of AgNWs 11 cycles, AgNWs 20 cycles films and of AgNWs 20 cycles with IZO 140 nm film. Bottom image shows a cross section of the film; (d) Transmittance at 550 nm versus sheet resistance for various materials - IZO, AgNWs, AgNWs + IZO, and films from literature; (e) AFM images of AgNWs 20 cycles with IZO 140 nm and IZO 140 nm films, with IZO evidencing a smooth surface, typical of amorphous films; (f) Resistance variation of IZO and AgNWs films when immersed in saline solution at 37 °C to simulate physiological conditions (SPC).

The optical, electrical and morphological properties of the films were analyzed. The optical transmittance measurements were taken with a PerkinElmer Lambda 950 UV/VIS/NIR Spectrometer. The measurements were performed using an integrating sphere in the wavelength range from 200 nm to 1500 nm. Samples with a Van der Pauw structure were used in a Hall effect system (BioRad HL5500) with a 0.5 T permanent magnet to access their sheet resistance at room temperature. The thickness of IZO was measured by profilometry using an Ambios XP‐200 profilometer. The surface morphology was investigated using Scanning Electron Microscopy (SEM, Zeiss Auriga Crossbeam microscope) and Atomic Force Microscopy (AFM) analysis was performed using an Asylum Research MFP-3D Stand Alone AFM system. SEM images were also used to calculate the areal mass density (amd), assuming uniform NWs network. Topography measurements were performed in tapping (alternate contact) mode in air, using commercially available silicon AFM probes (Olympus AC160TS; k = 26 N/m; f_0_ = 300 kHz).

Next, we quantified the interference of these films during fluorescence microscopy. Biological specimens for imaging were obtained from a C57Bl6 mouse (Charles River Labs, UK) expressing mCherry sparsely in neurons of the Lateral Posterior nucleus (LP) of the thalamus and several areas of the neocortex. The mouse had previously been transcardially perfused under terminal general anaesthesia with 0.1 M phosphate-buffered solution (PBS), followed by 4% paraformaldehyde (PFA) in PBS and the brain dissected from the cadaver. After 24h post-fixation in 4% PFA, the brain had been embedded in 4% agarose and 40 μm-thick brain slices were cut using a vibratome. To visualise cell nuclei, slices were incubated in Hoechst solution (20 mM, ThermoFisher Scientific UK, 33342) for 15 minutes and then washed 4 times in 0.1M PBS. Slices were mounted on glass slides, air-dried, covered in ProLong Glass Antifade Mountant (ThermoFisher Scientific UK, P36980) and then cover-slipped. The cover-slips were coated with the films under study. Imaging was performed in a Zeiss AxioImager 2 microscope (Apotome optical sectioning mode, 0.61 μm optical sections) under a Plan-Neofluar 20x/0.5 NA objective (Carl Zeiss, Jena, Germany).

### Production and characterization of ECoG EAs

Following films characterization implantable neural EAs were fabricated. Firstly, silicon carrier wafers were coated with PVA (88% hydrolyzed, average M.W. 20,000-30,000, Acros Organics) to help the peel-off process latter. The solution of PVA (100 mL of H_2_O with 5 g of PVA) was prepared on a hot plate at 80° C, where PVA was slowly added to the water at 80° C and stirred at 500 rpm for approximatly4 hours. Then, PVA solution was spin-coated for 60 s at 1000 rpm with acceleration of 200 rpm/s. At the end, the silicon wafer was moved to a hot plate at 95° C for 5 minutes. In the second step of the process, a ~5 μm thin layer of Parylene C (poly(para-chloroxylylene, SCS - Specialty Coating Systems) was deposited using a Specialty Coating System Model 2010. In the third step, for the hybrid electrodes, AgNWs were deposited onto the Parylene substrate and patterned using shadow masking to form electrodes, connection pads and interconnect lines. These masks were fabricated from a ~0.4 mm thick aluminum sheet, where the pattern was cut by a Gravograph IS400 with the blade positioned at a 10° angle to the perpendicular surface. The AgNWs network was then coated with a 140 nm thick IZO film. The connection pads area was protected during IZO deposition. For the IZO-only electrodes a film of 220 nm was deposited and patterned using shadow masking. In the fourth step, the connection pads to zero insertion force (ZIF) connectors and initial portions of the interconnect lines (excluding the electrode area) were prepared with Ti (± 6 nm) and Au (± 60 nm) via Electron Beam Evaporation using shadow masking. Gold was used to reduce line resistance and to improve the contacts between the devices and the ZIF connectors, whilst titanium was used to improve adhesion. Next, a ~1 μm thick passivation/insulating layer of parylene C was deposited. Plasma etching (RIE Trion Phantom III) was then used for etching Parylene C passivation layer, simultaneously opening windows for the electrodes and contact/ZIF interfaces. Finally, the arrays were released from the silicon carrier wafer by scratching the borders of the carrier and immersing in de-ionized water at 90° C for around 1 h (PVA started to dissolve and Parylene C to detach from the silicon wafer). Prototypes were then characterized by connecting them to an adapter^18^ equipped with an Omnetics connector, which is compatible with all measuring equipment (see Figure 2 in supplementary information).

**Figure 2.**
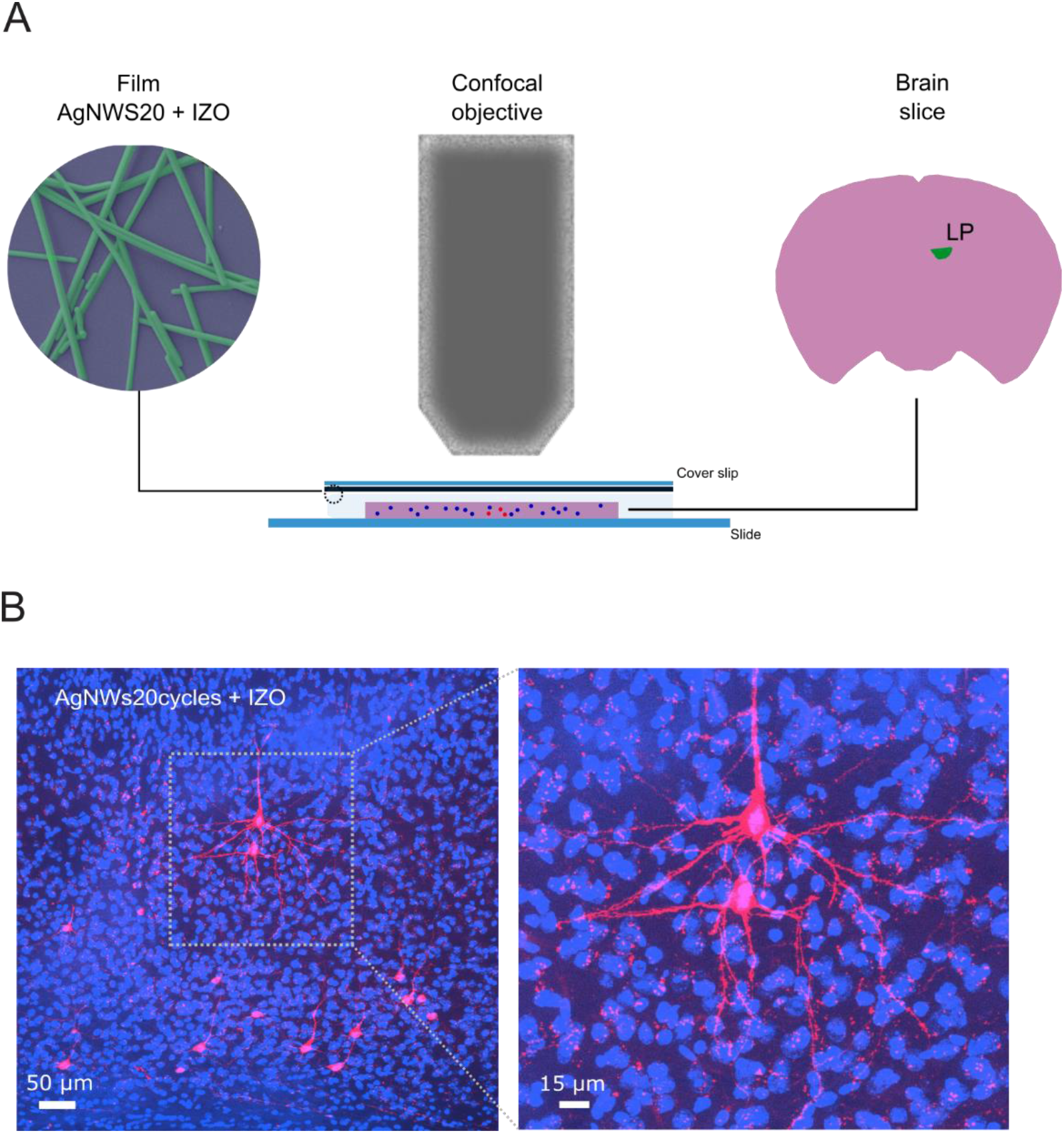
Optical imaging with fluorescence microscopy and hybrid films. (a) Schematic illustration of the confocal microscopy setup. A mouse brain slice was mounted on a glass slide and it was covered with a cover slip coated with the film under study. Note that excitation as well as emission light passes through films; (b) Multiples cells can be clearly imaged through the film.

The electrochemical behavior of the microelectrodes was studied in PBS 1 mM, pH 7.4 by electrochemical impedance spectroscopy and cyclic voltammetry. Electrodes were immersed in PBS and a NanoZ (Neuralynx) was used to characterize impedance, with a two electrode cell configuration - electrodes acted as working electrode and an Ag/AgCl wire as reference electrode (Science Products GmbH, E-255). Moreover, the impedance magnitude at 1 kHz of each electrode was also measured using a protocol implemented by the RHD2000 series chip (InTan Technologies). For the cyclic voltammetry, a potentiostat (Reference 600, Gamry Instruments) was used with a three electrode cell configuration where the electrodes were connected individually as the working electrode, a platinum wire served as the counter electrode, and an Ag-AgCl (3 M KCl, Gamry Instruments) as the reference electrode.

We measured the light-induced artifacts by immersing electrodes into PBS solution while using an Ag-AgCl wire as a reference (Science Products GmbH, E-255). We then used a fiber-coupled Light Emission Diode (LED) (LED 470 nm, fiber cable-SMA to 1.25 mm ferrule, and fiber optic cannula-200 μm, 0.39 NA, Thorlabs) to deliver blue light. By touching with the fiber tip on the microelectrode surface, artifacts were quantified from noise recording in the form of potential peaks. We studied the dependence of the amplitude of artifacts on different light intensity using a 10 ms pulse duration at 1 Hz. A power meter measured the light intensity used, which ranged from 14 to 95 mW/mm^2^. To drive the LED intensity and pulses a Pulse Pal^19^ and Bonsai workflow^20^ were used to enable precise sequences of voltage pulses. For the signal recordings we used the Open Ephys (http://www.open-ephys.org) acquisition board along with the RHD2000 series interface chip that amplifies and digitally multiplexes the signal from the 32 extracellular electrodes (Intan Technologies). Extracellular signals in a frequency band of 0.1-7,500 Hz were sampled at 20 or 30 kHz with 16-bit resolution and were saved in a raw binary format for subsequent offline analysis using Bonsai interface.

### Characterization of ECoG EAs in vivo – neural electrical recordings

The protocol was followed as described in previous works for urethane acute recording surgeries on rats^21^. A craniotomy was made over the left hemisphere, ~1-7 mm posterior and ~1 to 4 mm lateral to bregma. The transparent electrodes were positioned around 3 – 5 mm posterior relative to bregma, which in cortex covers motor cortex, primary somatosensory cortex, parietal cortex, and secondary visual cortex.

### Analysis

Python software was used to calculate the average peak-to-peak (P2P) amplitude of the light induced artifacts on a given electrode, where the analog signal from Pulse Pal was used for synchronization. Additionally, the magnitude of the background noise was estimated in saline solution as the standard deviation (root-mean-square noise) of the signal. Some results were represented as mean ± standard deviation. For the calculation of the power-spectral density (PSD) we used time frequency analysis tools from MNE package available at https://mne.tools/0.11/python_reference.html#time-frequency. We used epochs of 10 s and bandwidth of 0.5 Hz.

## RESULTS

### Characterization of transparent and conductive films

Figure 1 A and B illustrate transmittance spectra in the wavelength range from 250 to 1200 nm for IZO and hybrid films (AgNWs with IZO), respectively. Being transparent for optical imaging is important because it means that a large percentage of light is transmitted through the film in both directions, enabling excitation and emission light.

The measured thickness of both IZO films produced were 142 ± 17 nm (n = 4) and 225 ± 52 nm (n = 4). If we target for transmittance at 550 nm the IZO 140 and 220 nm films presented a value of 81.8 ± 0.5 % and 75.5 ± 1.1 %, respectively.

For the hybrid films, the transmittance at 550 nm decreased as more AgNWs are deposited. The network density varies with the number of deposition cycles. Above 7 deposition cycles the density is above the percolation threshold of the network, enabling a conductive path through the AgNWs that transports all charge carriers. Therefore, for the hybrid structure we selected AgNWs films with 11 and 20 cycles of deposition, which have densities above the percolation threshold (i.e., > 50 mg/m^2^). Figure 1 C shows the density of the nanowire networks achieved through 11 and 20 deposition cycles. As is evident from the SEM images in Figure 1 C, the AgNWs are randomly oriented and interconnected. Also, as shown in the same figure, the morphology of hybrid films, which are composed by AgNWs and IZO 140 nm coating, appeared identical to the uncoated ones, suggesting the IZO coating is covering perfectly the AgNWs network.

The AgNWs films with 11 and 20 cycles showed transmittances of 84.3 % and 76 % at 550 nm, respectively. The transmittance in the hybrid structure with AgNWs 11 cycles was 73.8 % and 61.7 % for the hybrid structure with AgNWs 20 cycles. Overall, AgNWs, IZO and hybrid films showed transmittances at 550 nm over 60 %, which is required for optical imaging^22^. Moreover, this transmittance does not decrease throughout the 400-1100 nm optical window. High transmittance in this broad frequency range enables different imaging techniques, such as widefield microscopy (~400-600 nm), two-photon microscopy (~800-1100 nm), and optogenetics (~470 nm) applications^12^.

The electrical properties of IZO, AgNWs and hybrid films were measured by Hall Effect. Figure 1 D summarizes transmittance values at 550 nm as a function of sheet resistance extracted from these films along with values reported in the literature for ITO^23,24^, graphene^25^ and patterned metal grids^12^. As expected, a tradeoff exists between conductivity and transparency. By increasing the thickness/density of material, the conductivity is increased but the transparency is decreased.

As shown in Figure 1 D the sheet resistance and the transmittance values achieved by our hybrid structures were similar to the metal grids^12^, whilst the IZO films presented sheet resistance and transmittance in the same range of values of ITO films used in transparent ECoG electrodes. Generally, our films when compared with other works, presented a lower sheet resistance. A low sheet resistance is important for fabricating interconnects that are long yet narrow. Moreover, from the plot in Figure 1 D we may conclude that the sheet resistance in hybrid structures is controlled by the AgNWs films and their sheet resistance varies with the network density. Nonetheless, the sheet resistance also relies on the nature of the interconnection between wires. The morphology of the hybrid film was additionally analyzed through AFM (Figure 1 E). AFM images are in good agreement with the SEM results. The IZO coating helps to reinforce the connectivity between the nanowires network, covering nanowires surface and filling the gaps between nanowires (increases the contact area between the wires). Nevertheless, IZO contribution to decrease the sheet resistance is small, less than 5%. Therefore, we conclude that the conductivity in the hybrid structure is mainly ensured by the AgNWs network due to efficient interconnections at wire junctions. More importantly, IZO coating improves AgNWs adhesion to the substrate (parylene C). The strong mechanical adhesion between the hybrid films to the substrate was confirmed via a Scotch tape test as described in^15^. The pristine AgNWs film was completely detached/erased from Parylene, whereas most film remains intact on the hybrid film (Figure 3 supplementary information). Furthermore, investigations on chemical stability (e.g., oxidation) of films for a period of approximately 30 days under simulated physiological conditions (37 °C and immersion in NaCl 0.9% solution) were conducted. As shown in Figure 1 F, the IZO film shows excellent chemical stability, with negligible variation of sheet resistance during this experiment, while the pristine AgNWs film presents a 5x increase after 34 days in the saline solution. Moreover, the change on sheet resistance for IZO and AgNWs films after being stored one year are plotted in Figure 1 F. No variation is seen for IZO, while the AgNWs films see its sheet resistance increased by 50 % from the initial value. All this characterization unveils the critical role of IZO to stabilize the AgNWs network for subsequent fabrication of reliable ECoG electrodes.

**Figure 3.**
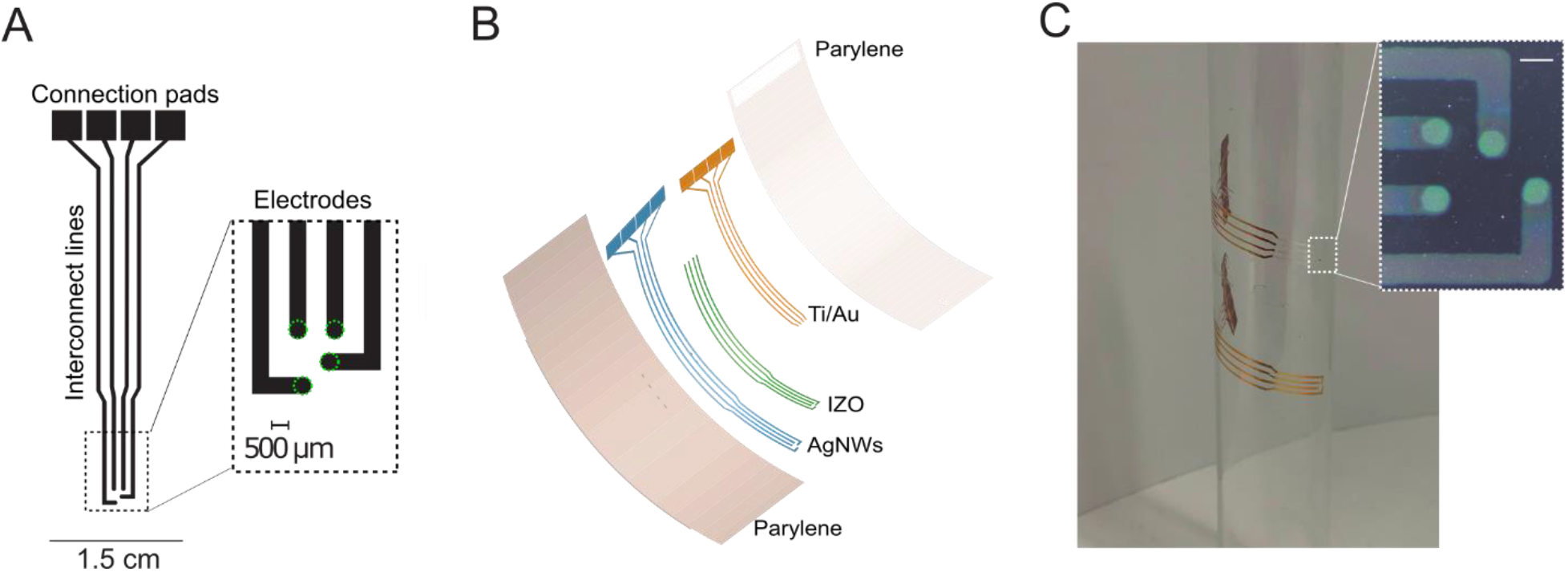
Electrode fabrication and overview. (a) Design of ECoG EAs prototype, including connection pads, interconnect lines and electrodes. The inset shows a magnified view of the EA, consisting of 2 × 2 electrodes (diameter 500 μm); (b) Diagram of prototype construction with hybrid films showing the flexible layered structure; (c) Demonstration of device flexibility. Prototypes are wrapped around a glass tub with a diameter of 2.5 cm. A gold prototype was also tested as control. Inset: optical microscope image of 4 electrodes. Scale bar: 500 μm.

Next, we enquired to which extent the presence of the films with the highest network density causes distortion of light during fluorescence microscopy. Firstly, we coated cover slips with the AgNWs20 + IZO film and then covered slipped brain slices expressing mCherry (excitation ~ 550 nm and emission ~ 600 nm) in neurons of the LP of the thalamus and several areas of the neocortex, and fluorescent dye (excitation ~ 358 nm and emission ~ 461 nm) in all neurons (Figure 2 A). Confocal microscopy images (Figure 2 B) suggest that hybrid AgNWs20 + IZO films have no impact on the optical imaging. Intensity profiles of neurons somas, across x and y axis, and along Z axis were plotted to quantify the full width at half-maximum (FWHM) obtained through hybrid films (Figure 4 supplementary information). The FWHM values for somas imaged trough glass (control) and hybrid films were similar in all axes (0- to 1.6-fold difference).

**Figure 4.**
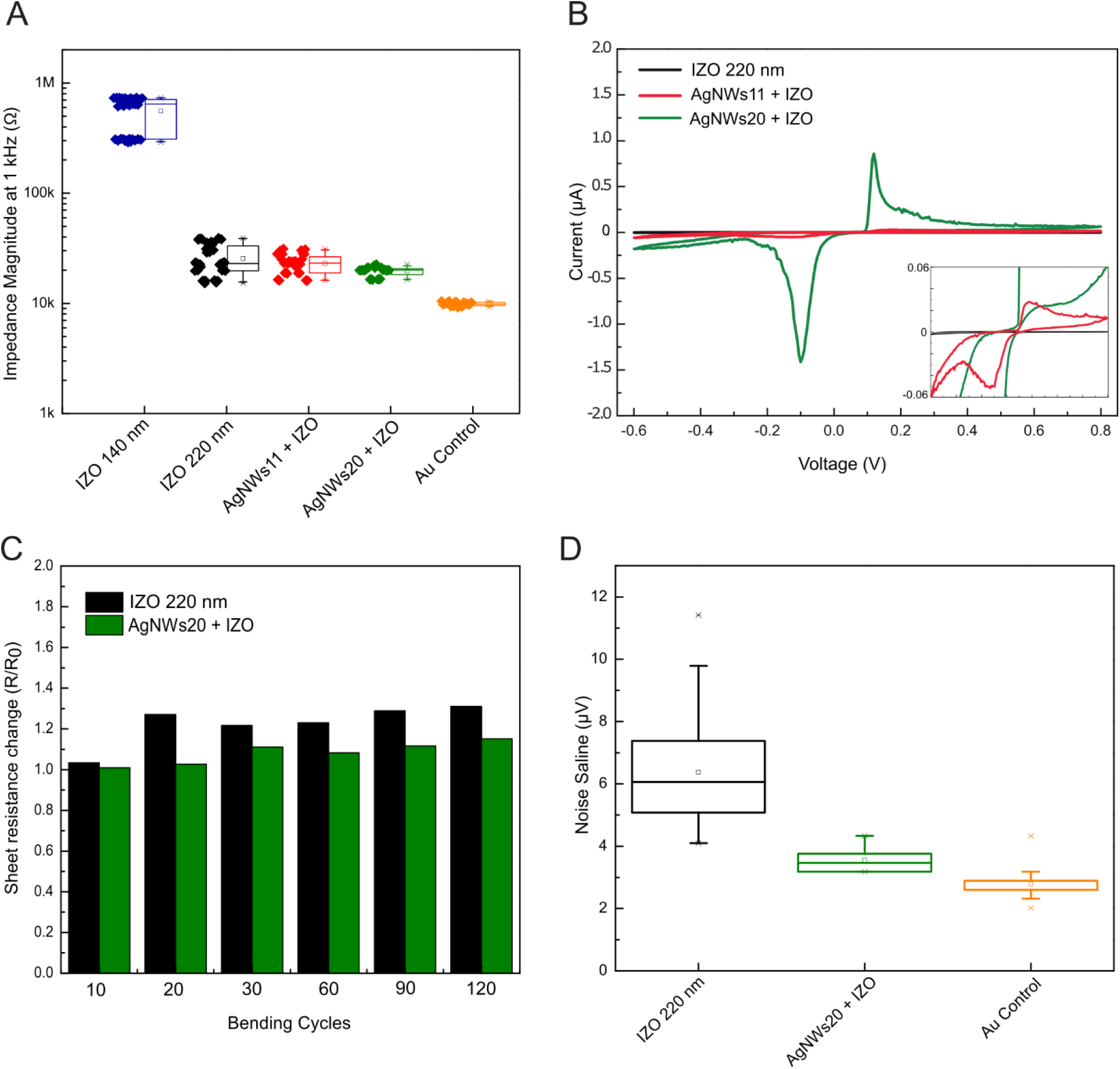
Electrodes characterization. (a) Impedance magnitude measured at 1 kHz in saline as a function of different electrodes and prototypes. Points denote impedance magnitude measured at 1 kHz in saline solution for individual electrodes (3 measurements for each electrode) and boxplots show the distribution of these values; (b) Cyclic voltammetry (3rd curve); (c) Mechanical characterization by testing resistance variation of IZO and AgNWs films during bending tests; (d) Root-mean-square noise of recordings (0.1–7.5 kHz) conducted in saline solution. In the boxplots, line: median, square: mean, box: 1st quartile–3rd quartile, and whiskers: 1.5× interquartile range above and below the box.

### Characterization of transparent and conductive ECoG EAs

Figure 3 A and B illustrate a schematic of the ECoG EAs and a schematic of the fabrication layers of a prototype with - Parylene C as substrate (5 μm thick) and encapsulation (1.5 μm thick) layer, Ti/Au as the contact pads and initial portions of the interconnect lines (6/60 nm thick), and AgNWs (11 or 20 cycles of deposition) and IZO (140 nm tick) as the electrodes and portions of interconnect lines. The final prototype is mechanically flexible due to its final thickness, less than ~ 7 μm (Figure 3 C).

Figure 4 shows the detailed electrochemical characterization of the electrodes. The impedance magnitude at 1 kHz is a common metric for assessing the performance of microelectrodes (Figure 4 A). The impedance of IZO 140 nm, IZO 220 nm, AgNWs11 + IZO, AgNWs20 + IZO at 1 kHz was 558 ± 185 kΩ, 25 ± 8 kΩ, 23 ± 5 kΩ, and 20 ± 2 kΩ, respectively. The impedance values at 1 kHz for the electrodes produced with IZO 220 nm and hybrid structures presented the lower impedance values. For further analysis, the cyclic voltammetry (CV) results of IZO 220 nm and hybrid structures appear in Figure 4 B. The enclosed area of CV curves defines the charge storage capacity (CSC) of the electrodes. From Figure 4 B, it is clear that the CSC value increases significantly with AgNWs density in the hybrid structure. The CSC derived from curves in Figure 4 B is 0.01 mC/cm^2^ for IZO 220 nm electrodes, 0.5 mC/cm^2^ for AgNWs11 + IZO electrodes and 3.2 mC/cm^2^ for AgNWs20 + IZO electrodes. The CSC value for AgNWs20 + IZO electrodes is higher when compared with the CSC value for the metal grids electrodes^13^. The increase in the superficial area, due to the increase in the AgNWs density, may explain the difference of CSC values between electrodes. As shown previously in Figure 1 E, the roughness of the surface and the superficial area increases in the hybrid structure when comparing with the IZO 140 nm film. Increased CSC for hybrid structures could enhance the amount of transferred charge when electrodes are used for neural stimulation. Furthermore, judging from the electrochemical impedance spectroscopy (EIS) measurements performed on electrodes, in the plot of phase as a function of frequency, as shown in Figure 5 supplementary information, the impedance became more capacitive (phase closer to 90°) due to the presence of AgNWs in the film. The overall impedance decreased as the electrode surface became rougher, probably due to the increased effective electrode surface area with enhanced charge transfer^26^. In conclusion, the hybrid AgNWs20 + IZO electrodes, which have the higher AgNWs density, have the best electrochemical performance.

**Figure 5.**
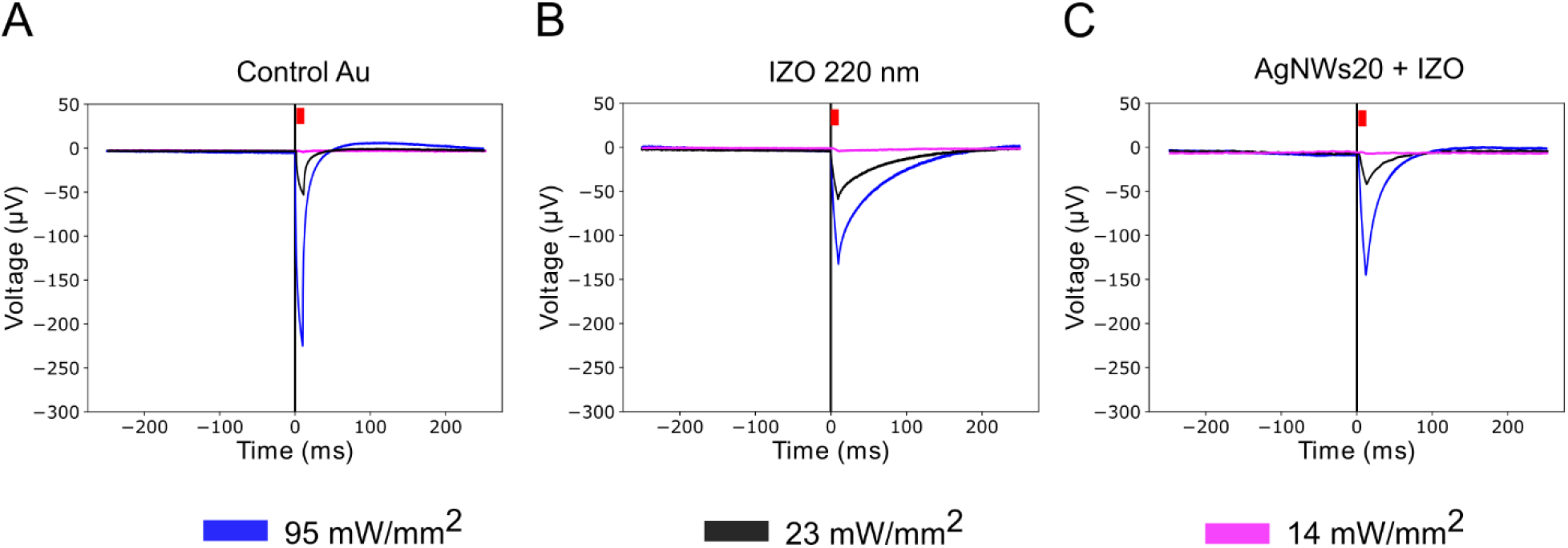
Light-induced artifact characterization in saline solution. (a-c) Averaged light-induced artifact of Control Au, IZO 220 nm and AgNWs20+IZO electrodes by varying light intensity. In each measurement a 10-ms light pulse was applied at 1 Hz for 1 min. The red colored trace represents the light pulse of 10 ms.

Mechanical flexibility and robustness of flexible ECoG EAs is also of great importance. In this work, we performed bending tests with a bending diameter of ~5 mm being the ratio of the change in film resistance recorded as a function of the bending cycle, as shown in Figure 4 C. The hybrid film has capability to endure mechanical bending showing a stable electrical performance, even after more than 100 bending tests. The presence of AgNWs network improves mechanical robustness of hybrid films. Nevertheless, this bending test represents an extreme situation for these films. For instance, during recordings from cortex surface, the curvature is smaller (larger bending diameter) and electrodes are implanted in a fixed position.

We also measured the noise in saline solution for the different electrodes, as shown in Figure 4 D. Root-mean-square (RMS) noise calculated from IZO, AgNWs20 + IZO and Au electrodes was 6.4± 0.3 μV, 3.5±0.1 μV, and 2.8±0.1 μV respectively. The noise measured in saline for gold and hybrid electrodes was thus similar and almost half of the recorded for IZO.

The presence of light-induced artifacts was quantified during electrical recordings in saline solution while applying optogenetic excitation light (470 nm). Results are shown in Figure 5 A-C including artifacts measured from Au, IZO 220 nm and AgNWs20 + IZO electrodes.

As expected, the artifacts increased with higher light intensity due to stronger photoelectric effects. For example, using 23 mW/mm^2^ light, the artifacts (black lines in Figure 5) peak-to-peak amplitude measured from Au microelectrodes was ~53 μV while those from IZO 220 nm and AgNWs20+IZO microelectrodes was about 63 and 38 μV, respectively. Our electrodes when compared with the metal grids electrodes^27^ have shown smaller artifacts for a given light intensity (e.g., the light-induced artifact at metal grids using 20 mW/mm^2^ was around 100 μV). Additionally, the peak-to-peak artifact amplitude using 14 mW/mm^2^ is within RMS noise value measured in saline solution, ~4 μV. Thus, for hybrid electrodes the light-induced artifacts shouldn’t be an issue when used in combination with functional calcium imaging -widefield and 2-photon microscopy- where the illumination intensity^27^ required is around 0.28 mW/mm^2^. Moreover, for optogenetic stimulation of neurons where the power is higher, around 100 mW/mm^2^, the light-evoked transient can be reduced by subtraction of the median of the signal across all channels^28^.

In Table 1, we summarized the transmittance, sheet resistance, impedance and CSC of IZO 200 nm and hybrid AgNWs + IZO electrodes to illustrate the trade-off between those parameters. We also benchmarked the performance of the fabricated electrodes against previous transparent electrodes. Our electrodes demonstrated comparable transmittance and impedance values, with improvements in the sheet resistance and CSC.

**Table 1.** Current transparent electrodes and their material, dimensions, transmittance value at 550 nm, sheet resistance, impedance magnitude at 1 kHz in saline solution, and charge storage capacity.

| Work | Material | Dimension | Transmittance @ 550 nm (%) | Sheet resistance ($\Omega/\text{sq}$) | $\vert Z\vert $ @ 1 kHz (kΩ) | CSC (mC/cm<sup>2</sup>) |
| --- | --- | --- | --- | --- | --- | --- |
| Kuzum et al., 2014 | Doped Graphene | 50x50 $\mu\text{m}$ | 90 | N.A. | ~500 | 1.95 |
| Ledochowitsch et al., 2011 and 2015 | ITO | 500 $\mu\text{m}$ $\oslash$ | 90 | 43.7 | 300 | N.A. |
| Park et al., 2014 | Graphene | 200 $\mu\text{m}$ $\oslash$ | ~90 | 76 | 243.5 | N.A. |
| Thunemann et al., 2018 | Graphene | 100x100 $\mu\text{m}$ | 90 | N.A. | > 963 | N.A. |
| Kunori & Takashima, 2015b | ITO | 250 $\mu\text{m}$ $\oslash$ | 80 | 200 | 10-50 | N.A. |
| Seo et al., 2017 | Au grid | 80x80 $\mu\text{m}$ | 70 | 18 | 127.2 | N.A. |
| Qiang et al., 2017 | Au grid+ PEDOT | 80x80 $\mu\text{m}$ | 70 | N.A. | 11 | 2.55 |
| Lu et al., 2018 | Graphene<br>30 sec | 100x100<br>$\mu\text{m}$ | 50 | N.A. | 15 | N.A. |
| This work | IZO 100 nm | 500 $\mu\text{m}$ $\oslash$ | 82 | 53 | 558 | N.A. |
| | IZO 200 nm | 500 $\mu\text{m}$ $\oslash$ | 76 | 25 | 25 | 0.01 |
| | AgNWs11+<br>IZO | 500 $\mu\text{m}$ $\oslash$ | 74 | 22.5 | 23 | 0.5 |
| | AgNWS20<br>+IZO | 500 $\mu\text{m}$ $\oslash$ | 62 | 5.9 | 20 | 3.2 |

$\oslash$ : diameter; N.A.: not available

### Characterization of ECoG EAs in vivo - electrical recordings

We next recorded neural activity from a rat cortex surface using transparent ECoG EAs fabricated with AgNWs20 + IZO hybrid films. Figure 6 A-B shows the position of electrodes and craniotomy. Figure 6 B demonstrates the clarity of the electrodes and interconnect lines and the ability to view the underlying cortex and cerebral vasculature through the prototype. Cortical electrical potentials recorded by our electrode array are shown in Figure 6 C. The electrical signal shows spontaneous cortical activity recorded on all four electrodes under urethane anaesthesia without any sensory stimulus. Urethane anesthesia induces sleep-like rhythms characterized by low frequency oscillations between 0.5 to 4 Hz (i.e. delta waves), which reflects a synchronized brain activity ^30–32^. We compared the ECoG signal recorded by our electrodes and the local field potential (LFP) signal recorded by a small size electrode (15 μm diameter) inserted 1245 μm deep into cortex under urethane anesthesia^33^. This dataset of recordings from anesthetized rats performed with polytrodes containing 32 electrodes at different cortex depths is freely available online (https://www.kampff-lab.org/polytrode-impedance). Power spectral density analysis (Figure 6 D) of both ECoG and LFP signals in the 1–100 Hz frequency range reveals higher power in low-frequency oscillations corresponding to delta band (i.e., 0.5 to 4 Hz), which is concomitant with sleep-like rhythms. This suggests that our ECoG electrodes are able to pick up the signals that have been found by intracortical electrodes under urethane anesthesia.

**Figure 6.**
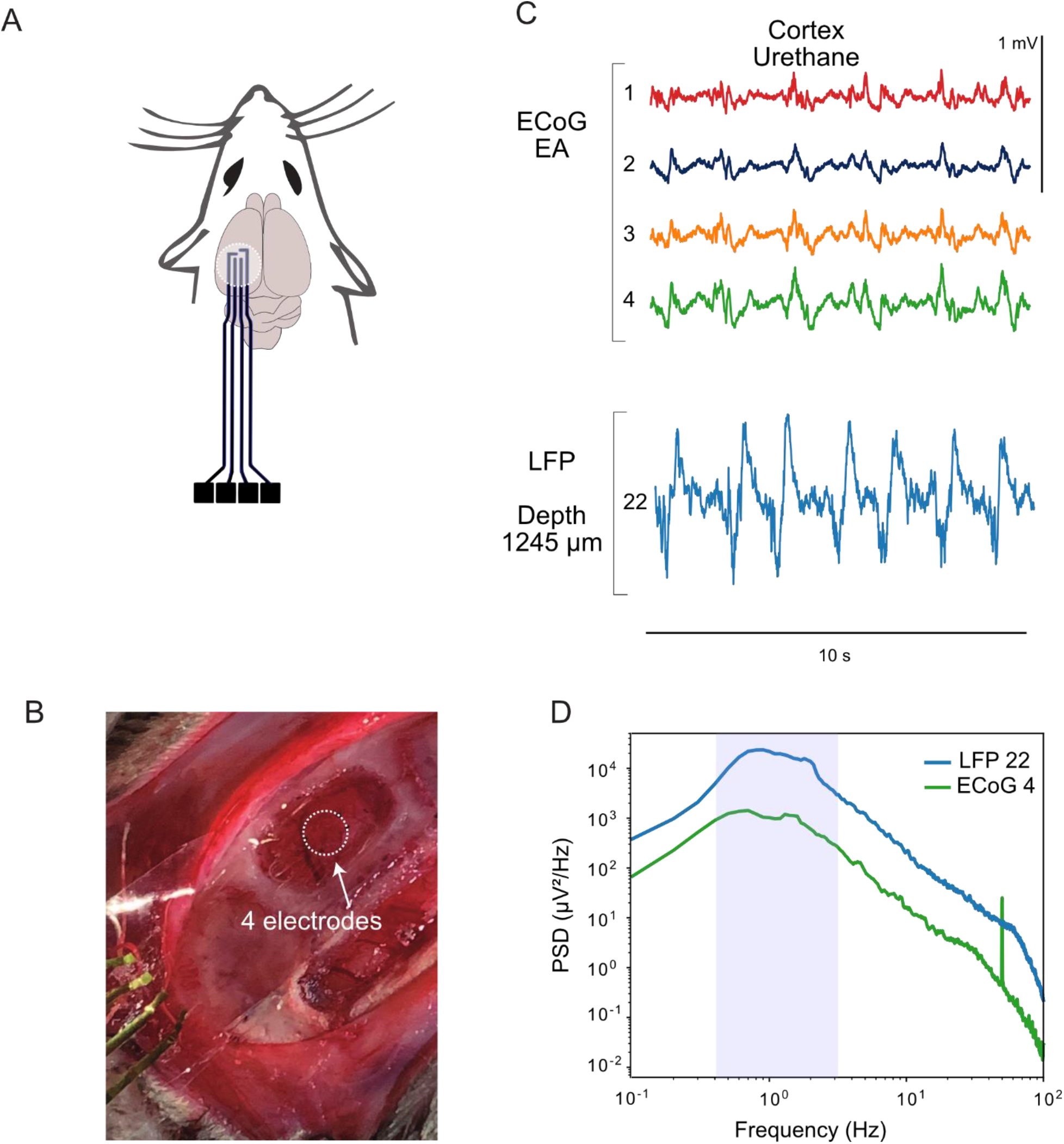
Characterization of electrodes in vivo. (a) Scheme of acute recordings in vivo, showing ECoG EA position; (b) Photograph of transparent and flexible ECoG EA fabricated with AgNWs20 + IZO films placed over the cortex of an anesthetized rat; (c) 10-s-long traces of ECoG electrodes and a depth electrode measuring the LFP in motor cortex under urethane anesthesia. The LFP signal corresponds to the electrode number 22 from recording amplifier2017-02-02T15_49_35.bin, whereas the distance (depth) from the brain surface to the electrode was 1245 μm. Both ECoG and LFP signals were low-pass filtered; (d) Signal power spectra density for one ECoG electrode (number 4) and one intracortical electrode (number 22) measuring neuronal electrical activity under urethane anesthesia.

## CONCLUSIONS

We demonstrate a new transparent material for flexible and stable ECoG electrodes with low sheet resistance, low impedance, high CSC and good transmittance.

The hybrid electrodes formed by depositing IZO onto random networks of AgNWs demonstrated transparency over a broad optical window (400 to 1200 nm), with over 60% transmittance at 550 nm, which is suitable for optical imaging measurements. Moreover, the films produced with the hybrid material show the lowest sheet resistance achieved among transparent ECoG electrodes reported in literature, as can be verified in Table 1.

These electrodes are successfully demonstrated for in-vivo electrical measurements, by recording neural activity from a rat cortex, being able to capture the signals associated with sleep-like rhythms.

The results presented here can be further enhanced, particularly in terms of optical transmittance, by simply replacing the thick IZO sputtered films by ultra-thin (i.e., few tens of nm), conformable and conductive oxide coatings made by industrial-compatible techniques as atomic layer deposition (ALD). By introducing ALD-based indium-free materials environmental sustainability and overall process cost can also become even more advantageous over competing ECoG electrode technologies.

## ASSOCIATED CONTENT

### Supporting Information

Scheme of one cycle of AgNWs spray deposition; photograph of adapter with a ZIF and Omnetics connector; results of adhesion test using Scotch tape; full width at half-maximum (FWHM) obtained from intensity profiles of neurons somas through hybrid films during fluorescence microscopy; EIS measurements on prototypes with different films.

## AUTHOR INFORMATION

## ACKNOWLEDGMENT

We would like to thank the institutional support and funding provided by Sainsbury Wellcome Centre (funded by the Gatsby Charitable Foundation and the Wellcome Trust).

This work was funded by FEDER funds through the COMPETE 2020 Programme and National Funds through the FCT – Fundação para a Ciência e a Tecnologia, I.P., under the scope of the project UIDB/50025/2020. This work also received funding from the European Community’s H2020 program under grant agreement No. 716510 (ERC-2016-StG TREND).

We are very grateful to George Dimitriadis and Lorenza Calcaterra for animal experiment support. We also thank Tiago Mateus, Ana Santa, Inês Martins, Cátia Figueiredo, Tomás Calmeiro and Daniela Gomes for technical support.

## ABBREVIATIONS

AgNWs: silver nanowires
ECoG: electrocorticography
EA: electrode array
AFM: atomic force microscopy
LP: lateral posterior nucleus
EIS: electrochemical impedance spectroscopy
ZIF: zero insertion force
SEM: scanning electron microscopy
amd: areal mass density
PBS: phosphate-buffered solution
P2P: peak-to-peak
RMS: root-mean-square
PSD: power-spectral density
SPC: simulated physiological conditions
FWHM: full width at half-maximum
CV: cyclic voltammetry
CSC: charge storage capacity
LFP: local field potential
ALD: atomic layer deposition.

## Notes

### Competing Interest Statement

The authors have declared no competing interest.

### Summary of Updates

In this second version experimental work was conducted with the aim of understanding IZO contribution in hybrid electrodes stability and performance.

