## supplementary information for "Transparent and flexible ECoG electrode arrays based on silver nanowire networks for neural recordings"

**List of figures:**

**Figure 1.** Scheme of one cycle of AgNWs spray deposition.

**Figure 2.** Photograph of adapter with a ZIF and Omnetics connector. Prototypes are connected via ZIF connector (FLZ connector, 36FLZ-RSM1-R-TB). This PCB equipped with Omnetics connector is compatible with NanoZ and Intan Headstage.

**Figure 3**. Optical microscope images of the AgNWs 11 and the hybrid (AgNWs 11 + IZO) films on the substrate after the adhesion test using Scotch tape.

**Figure 4.** Quantification of full width at half-maximum of neurons expressing mCherry (red) and dye (blue) imaged trough glass (control) and hybrid films (AgNWs20 + IZO). The intensity profiles of neurons somas were plotted across x and y axis and along Z axis using Image J (plot XY profile and plot Z-axis profile within a rectangular selection surrounding soma). Inset shows a representative intensity profile in Z axis from a neuron expressing mCherry. Mean ± standard deviation per condition are shown.

**Figure 5.** EIS of prototypes with IZO 220 nm, hybrid films and Au electrodes.

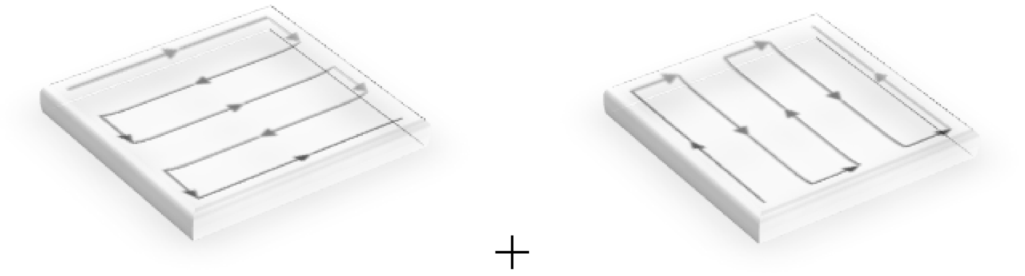

**Figure 1.** Scheme of one cycle of AgNWs spray deposition.

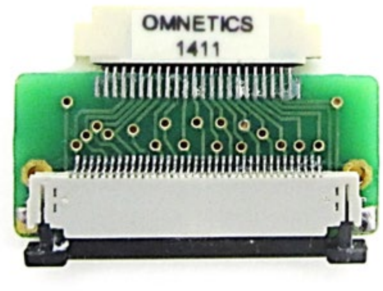


**Figure 2.** Photograph of adapter with a ZIF and Omnetics connector. Prototypes are connected via ZIF connector (FLZ connector, 36FLZ-RSM1-R-TB). This PCB equipped with Omnetics connector is compatible with NanoZ and Intan Headstage.

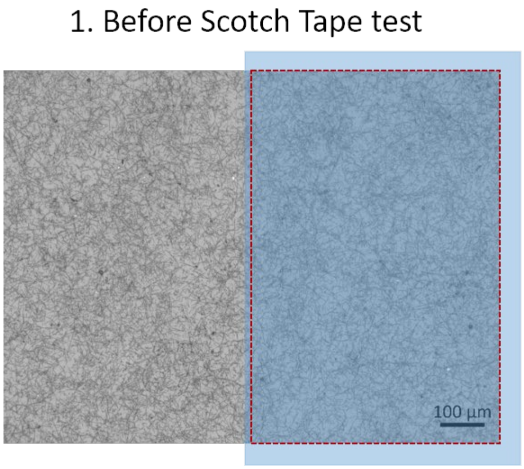


**Figure 3.** Optical microscope images of the AgNWs 11 and the hybrid (AgNWs 11 + IZO) films on the substrate after the adhesion test using Scotch tape.

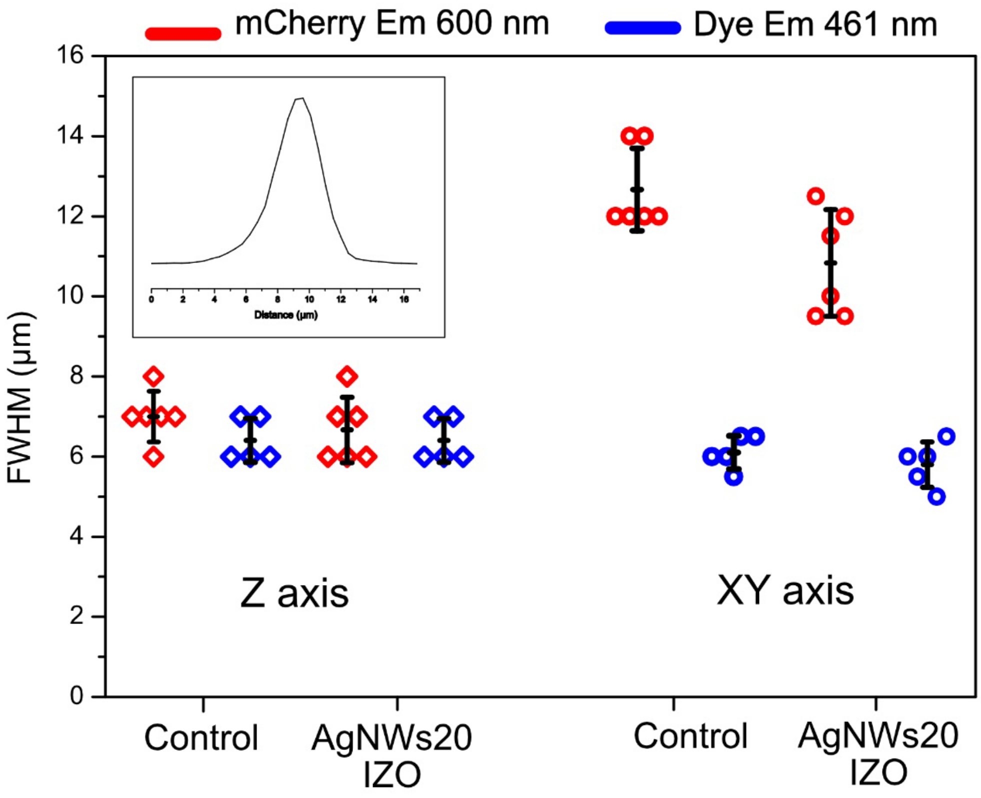


**Figure 4.** Quantification of full width at half-maximum (FWHM) of neurons expressing mCherry (red) and dye (blue) imaged trough glass (control) and hybrid films (AgNWs20 + IZO). The intensity profiles of neurons somas were plotted across x and y axis and along Z axis using Image J (plot XY profile and

plot Z-axis profile within a rectangular selection surrounding soma). Inset shows a representative intensity profile in Z axis from a neuron expressing mCherry. Mean ± standard deviation per condition are shown.

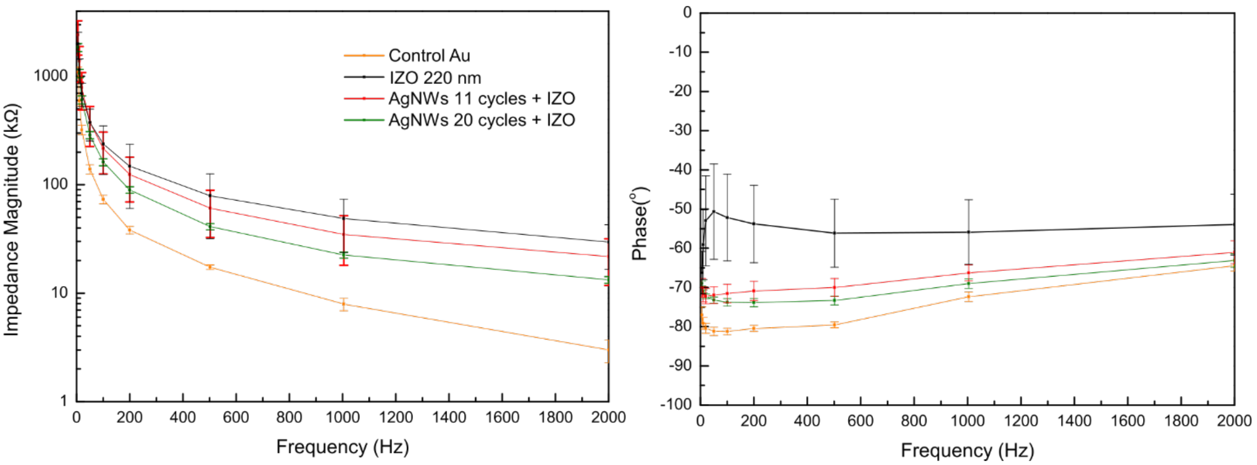


**Figure 5.** EIS of prototypes with IZO 220 nm, hybrid films and Au electrodes.
